# Assessing the haplotype and spectro-functional traits interactions to explore the intraspecific diversity of common reed in Central Italy

**DOI:** 10.1101/2022.12.19.521025

**Authors:** Maria Beatrice Castellani, Andrea Coppi, Rossano Bolpagni, Daniela Gigante, Lorenzo Lastrucci, Lara Reale, Paolo Villa

## Abstract

As reflectance measured via remote sensing is connected to plant light use and morpho-structural features, it can be used to derive spectral proxies of functional traits, or spectro-functional traits. Focusing on disentangling intraspecific trait variability in nature, we evaluated the links between haplotype and spectro-functional traits in *Phragmites australis* populations.

Haplotypes sequencing and multi-seasonal satellite data were used to evaluate the temporal dynamics of spectro-functional traits for reed stands sampled from seven wetlands in Central Italy, investigating meteo-climatic drivers, the differences across ecological statuses, sites, and haplotypes, and quantifying intraspecific variability due to haplotype or phenotypic plasticity.

Five haplotypes were identified, including an unedited one, which explained a substantial portion of intraspecific variability in canopy traits, differing for aquatic and terrestrial stands. We found that meteo-climatic factors impact on aquatic reeds traits (not over terrestrial ones) and a dualism between most and less common haplotypes, pointing to different evolutionary strategies. Dynamics in reed canopy traits were linked to ecological status, site and haplotype, with signs of haplotype-variable effects of dieback on aquatic stands.

Evaluating the spectro-functional variability over reed haplotypes may provide a straightforward approach for monitoring the genotype-phenotype relations across scales and assessing their ecological drivers.

## 1. Introduction

Because of their sessile condition, plant species are largely prone to respond to changes in environmental conditions by varying multiple traits (Valladares et al. 2007). Even if the basic assumption underlying comparative ecology studies is that functions are more variable among than within species (McGill et al., 2006), recent studies focusing on intraspecific trait variability (ITV) have shown that this does not hold for all the traits of a species in equal ways (Albert et al., 2010).

Different environmental filters may drive the selection of different genotypes and phenotypes in the context of population assembly rules, with abiotic and biotic factors tending to act idiosyncratically (Violle et al., 2012; Garnier et al., 2016). Given the strong environmental filtering acting on aquatic and wetland habitats, species replacement is more difficult than in terrestrial ones and aquatic plants are generally showing a high degree of intraspecific variability to cope with their habitat changes (i.e., see Jung et al., 2010 regarding the variability of SLA along a flooding gradient). In highly dynamic wetland ecosystems, not only abiotic factors (e.g., water level, nutrients availability) can be extremely variable even at local scales, but also biotic factors (e.g., herbivore pressure, competition with phytoplankton, sediment microbiome composition) can show fine scale heterogeneity in spatial and temporal patterns, all together acting on performance and selection of aquatic vegetation communities (Yang et al., 2019; Stagg et al., 2020).

Within this frame, the geographic distribution of a species plays a role too, as it is widely accomplished that species broadly distributed tend to be more variable due to local adaptation and/or acclimation across a broad range of environmental conditions (Eller et al., 2017; Pither, 2003; Siefert et al., 2015). This intraspecific variation enables widespread species to have broad ecological amplitudes in response to biotic (e.g., herbivores, mutualists, pathogens) and abiotic (e.g., climatic, edaphic) environmental factors (Albert et al., 2010; Baythavong & Stanton, 2010).

In this respect, *Phragmites australis* (Cav.) Trin. ex Steud. is one of the most distributed of all flowering plants, showing a broad ecological amplitude and intraspecific variability (Meyerson et al., 2016, Coppi et al 2018). This species grows in very diverse wetland habitat types, such as riverbanks, lakeshores, brackish and salty marshes, in oligotrophic to eutrophic waters and soils (Landucci et al., 2013). In these habitats, *P. australis* has important ecological, economic, and social roles (Kiviat, 2013; Ostendorp, 1993), and forms large and monospecific stands (Granéli, 1989) that, in some cases, displace other wetland plants (Foggi et al., 2011; Próchnicki, 2005). Thanks to the analysis of two noncoding chloroplast regions, Saltonstall (2002) described a total of 27 haplotypes from worldwide specimens of *P. australis* and depicted a cryptic invasion process by a non-native genotype of common reed in North America. Subsequently Lambertini et al. (2012), based on analysis of chloroplast variation with parsimony and genetic distance methods, extended the knowledge on haplotype occurrence in natural populations of European *P. australis*, adding four novel haplotypes detected in Romania and Northern Europe. More recently, Coppi et al. (2018) showed the occurrence of five different haplotypes in central Italy (M, K, CO, VI, and CHTR) mostly forming patches in the same wetland, and Naugžemys et al. (2021) identified six new haplotypes (BM, BN, BO, BP, BQ and BR) in Lithuania river sites, alongside two known European haplotypes (M and L).

Partitioning of intraspecific variability of regional populations (e.g., few genotypes and highly plastic *vs* many genotypes with low plasticity) is essential for assessing adaptation potential over time scales, also in face environmental change (Benito Garzón et al., 2011; King et al., 2018). Within this framework, there is a need for tools for assessing intraspecific plant diversity across scales, which are both reliable and efficient (Valladares et al., 2006). For its synoptic capabilities and the availability of abundant operational satellite constellation put on orbit in the last decade, the exploitation of remote sensing data for plant functional studies is nowadays a strong option (Jetz et al., 2016) - in particular towards the monitoring of plant canopy morpho-structural and biochemical characteristics and their spatial and temporal dynamics (see for example the reviews of Homolová et al., 2013; Verrelst et al., 2015; Gamon et al., 2019; Dalla Vecchia et al., 2020).

Remote sensing platforms acquiring data in the visible to shortwave infrared spectral range (400-2500 nm), the so-called passive optical systems, are used to measure vegetation spectral reflectance, a physical quantity (directly or indirectly) connected to light use and photosynthesis, and can be therefore used for characterizing plant functional types (e.g., Ustin & Gamon et al., 2010; Villa et al., 2015; Schweiger et al., 2017), as well as for modelling functional traits at leaf to canopy scales (e.g., Asner & Martin, 2008; Asner et al., 2015; Serbin et al., 2019). Due to their straightforwardness, spectral indices (SIs) derived from optical remote sensing data as algebraic combinations of reflectance in different spectral bands, can be used as proxies of specific functional traits: e.g., canopy density, biomass, leaf area index (e.g., Le Maire et al., 2008; Ustin et al., 2009; Villa et al., 2017; Feilhauer et al., 2018; Van Cleemput et al., 2019). As this, SIs could be easily employed within a framework for assessing intraspecific diversity in monospecific or quasi-monospecific communities, where satellite data pixels (usually metric to decametric resolution) are almost completely comprising one single species canopy. In similar plant community setup, such is typically the case with *P. australis* stands, decametric pixels of multispectral satellite scenes provide an unambiguous link to spectral response of the individual species in its natural environment at canopy scale, and satellite derived plant functional proxies can be used for assessing intraspecific diversity across sites and seasons.

The main aim of this study was to explore the links between haplotypes and spectral proxies of canopy functional traits (i.e. spectro-functional traits) in natural stands of *P. australis* from Central Italy, taking into account meteo-climatic and environmental conditions (site characteristics and ecological status of stands). The evaluation of spectro-functional variability over genotyped common reed patches should indeed represent a straightforward and efficient approach for monitoring the haplotype-phenotype relations across geographical and temporal scales, and eventually contextualize them in terms of environmental drivers and ecological significance.

## 2. Methods

### 2.1. Study area and plant materials collection

The study areas are seven wetlands in central Italy (Figure 1, Table 1), including two marshlands (Fucecchio and Colfiorito) and five lakes (Chiusi, Massaciuccoli, Porta, Trasimeno, and Vico). The seven sites differ in their pedo-morphological characteristics, passing from lowland and upland wetlands such as Colfiorito, Fucecchio, Massaciuccoli, and Porta to shallow, turbid lakes such as Chiusi and Trasimeno, or a deep, clear lake of volcanic origins as Vico. Surface areas are furthermore variable, with the smallest site (Colfiorito) covering 0.8 km^2^ to the largest one (Trasimeno) covering 121.5 km^2^. Leaf tissue was collected from each of the sampled stands, to implement the set of haplotypes described in Coppi et al. (2018). Repeated sampling during the growing seasons of 2020 was performed within selected plots. The final dataset was used to confirm the occurrence of haplotypes across the studied areas, expand the knowledge on the haplotypes’ occurrence in other study areas (e.g. lakes Porta and Massaciuccoli), and identify spatial patches over which to extract spectro-functional traits from satellite data. The complete list of the specimens examined, their provenance and the GenBank information retrieved are given in Supplementary material S1.

**Figure 1.**
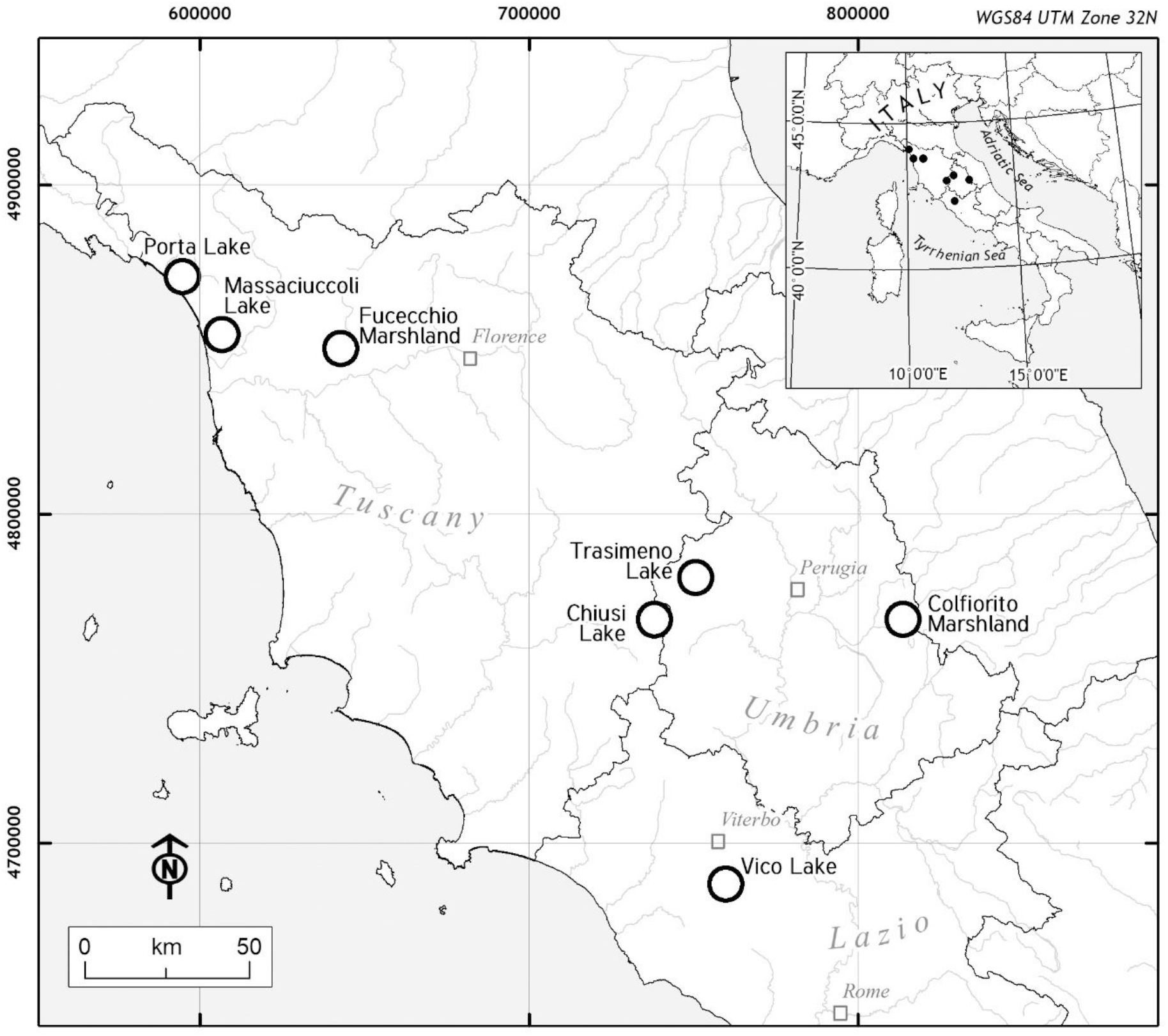
Geographic location of the study area with distribution of the sampling sites (empty dots).

**Table 1.** Overview of the geographical coordinates of the 7 study sites with locality, surface area, elevation and type of site information.

| Study site | Locality | Latitude | Longitude | Surface area (km<sup>2</sup>) | Elevation (m a.s.l) | Type of site |
| --- | --- | --- | --- | --- | --- | --- |
| Chiusi lake | South-eastern Tuscany | 43° 03' 22.11'' N | 11° 57' 55.79'' E | 3,87 | 245 | Shallow lake |
| Colfiorito marshland | Eastern Umbria | 43° 01' 23'' N | 12° 52' 36'' E | 0,8 | 750 | Palustrine wetland with small open-water areas |
| Fucecchio marshland | North-western Tuscany | 43° 48' 30.38'' N | 10° 48' 20.14'' E | 1,02 | 15 | Palustrine wetland with small open-water areas and canal nets |
| Massaciuccoli lake | North-western Tuscany | 43° 50' 34'' N | 10° 19' 21'' E | 6,9 | 0 | Shallow lake |
| Porta lake | North-western Tuscany | 43° 59' 27'' N | 10° 10' 23'' E | 1,558 | 0 | Palustrine wetland with small open-water areas |
| Trasimeno lake | North-western Umbria | 43° 08' 05.5'' N | 12° 06' 04.6'' E | 128 | 255 | Shallow lake |
| Vico lake | North-western Lazio | 42° 18' 58.40'' N | 12° 10' 5.89'' E | 12,93 | 510 | Deep lake |

*P. australis* stands were further classified following the model proposed by Lastrucci et al. (2016c) regarding canopy background conditions at the peak of summer in terms of soil moisture and submersion. The ecological status of sampled plots was summarized into three categories: flooded stands (F), when the rhizomes were found covered by water all the year; floating island stands (Fl), when plants were found forming mats with rhizomes floating above the water; and dry stands (D), when the surfaced of soil substrate showed different degrees of wetness and dried for at least one month in the hottest period of the year. Category Fl includes only some plots from Lake Chiusi, while categories F and D include plots from all surveyed sites.

### 2.2. DNA extraction and amplification

Genomic DNA was extracted from silica-gel-dried leaf samples using a modified 2× cetyltrimethylammonium bromide protocol GenElute Plant Genomic DNA Miniprep Kit (Sigma). Amplification of the trnT-trnL and rbcL-psaI intergenic spacer regions of the cpDNA was performed following the protocol described by Taberlet et al. (1991) and Saltonstall (2001, 2002). Automated DNA sequencing was performed directly from the purified PCR products, using BigDye Terminator version 2 chemistry, and a sequencer (ABI310; PE-Applied Biosystems, Norwalk, CT, USA). To reduce artefacts, all the samples were amplified and sequenced twice, using both forward and reverse primers.

### 2.3. Sequence alignment and data analysis

A total of 178 original sequences of *P. australis* (89 for each of trnT-trnL and rbcL-psaI markers) were checked using BioEdit ver. 7.0 (Hall et al., 1999) through comparisons with accessions published in Coppi et al. (2018) and retrieved from the National Center for Biotechnology Information (NCBI). A single dataset that comprised the concatenated sequences of the intergenic spacer regions (trnT-trnL+rbcL-psaI) underwent fitting and analysis. The haplotypes identification and tagging follow the model described by Saltonstall (2001, 2002), implemented more recently by Hauber et al. (2011), Lambertini et al. (2012), Saltonstall (2016) and Coppi et al. (2018). Variation in the number of mononucleotides in the trnT-TrnL and rbcL-psaI sequences, was attributed to the variation at intra-haplotypic level and, as reported in Lambertini et al. (2012) and by Saltonstall (2016), has not been used to define new haplotypes.

### 2.4. Meteo-climatic data

ERA5-Land monthly averaged data from model reanalysis (Muñoz Sabater, 2019), managed by ECMWF and Copernicus, with 0.1° resolution and covering the period from January 1981 top December 2020: air temperature at 2 m above the surface, accumulated total evaporation from Earth’s surface, accumulated total incoming solar (shortwave) radiation reaching the Earth’s surface.

### 2.5. Satellite data

Satellite data used are surface reflectance products (L2A) derived from Sentinel-2 constellation, jointly managed by the EU Copernicus programme and European Space Agency (ESA). The two satellites composing Sentinel-2 constellation, i.e. Sentinel-2A (operational since July 2015) and Sentinel-2B (operational since July 2017), carry on board the same multispectral camera: the MultiSpectral Instrument (MSI), a push-broom imaging sensor with 13 spectral bands covering visible to shortwave infrared range (440-2250 nm) at medium spatial resolution (10 to 60 m pixel side on the ground), with a revisit time (in cloud free conditions) of 10 days with one satellite and of 5 days with both satellites (Drusch et al., 2012).

Cloud-free Sentinel-2 scenes acquired over each study area in July to early August, i.e. representing reed stands at their seasonal peak of density and greenness, were gathered from the existing archives for the years 2015-2020. After a preliminary assessment of differences in overall eco-environmental conditions of reed stands falling within the sampled sties at the time of satellite data acquisitions, compared to the season when *in situ* data collection took place (in 2014, 2017 or 2020, depending on the sampled reed stand), some years were deemed as not suitable for the analysis, i.e. year 2018 for Colfiorito, years 2017, 2019 and 2020 for Fucecchio, and year 2017 for Massaciuccoli, and were therefore excluded.

The satellite dataset used for the analysis of intraspecific variability of reed populations in terms of their spectral functional proxies was finally composed of the Sentinel-2 scenes listed in the following:

- Chiusi: 11/07/2015; 18/07/2016; 15/07/2017; 15/07/2018; 20/07/2019; 29/07/2020;
- Colfiorito: 11/07/2015; 04/08/2016; 15/07/2017; 20/07/2019: 24/07/2020;
- Fucecchio: 04/07/2015; 18/07/2016; 18/07/2018;
- Trasimeno: 11/07/2015; 18/07/2016; 15/07/2017; 15/07/2018; 20/07/2019; 29/07/2020;
- Vico: 04/07/2015; 18/07/2016: 10/07/2017; 10/07/2018; 05/07/2019; 29/07/2020;
- Massaciuccoli: 04/07/2015; 18/07/2016; 18/07/2018; 18/07/2019; 27/07/2020;
- Porta: 18/07/2019.

### 2.6. Spectral proxies of functional traits

Spectral indices known for their sensitivity to vegetation conditions at canopy scale were derived from Sentinel-2 scenes at 10m resolution as surrogates of specific reed functional traits, or spectro-functional traits: i) the Water Adjusted Vegetation Index (WAVI), a proxy of aquatic vegetation canopy density and fractional cover (Villa et al., 2014); ii) the Green Leaf Index (GLI), a proxy of canopy greenness and fraction of absorbed PAR (Hunt et al., 2011); and iii) the Normalized Difference Spectral Index for LMA (NDSI_LMA_), a proxy of leaf mass per area in macrophytes, aggregated at canopy scale (Villa et al., 2021).

All values of the 3 spectral proxies falling within a polygon surrounding the plots of common reed communities sampled *in situ* (covering 3-13 pixels, equal to 300-1300 m^2^ area) were extracted and used for calculating the mean and coefficient of variation (CV) of WAVI, GLI and NDSI_LMA_.

Following this, matchups between Sentinel-2 derived spectro-functional traits for different years with *in situ* samplings were retained only when difference dates between satellite scene acquisition and in situ sample collection did not exceed 2 years. Moreover, some matchups were discarded in case of marked differences in eco-environmental conditions between the sampling date and satellite acquisition, i.e. due to ambiguities in plot recorded coordinates (one plot in Colfiorito and one in Trasimeno) or to strong inter-seasonal differences in water level (observed for Fucecchio and Trasimeno sites). Finally, polygons scoring strong internal variability in functional proxies for one year, i.e. CV higher than 0.5 in at least one of the spectral indices, were excluded from the final list of matchups (Table 2) as they were considered not consistently representing reed canopy features, but more a mixture of spectral targets.

**Table 2.** Spectral proxies – in situ samplings matchup table (F = flooded stand, Fl = floating island stand, D = dry stand), divided by reed haplotype.

| Site | Type of site | BS | K | L | M | VI | TOT |
| --- | --- | --- | --- | --- | --- | --- | --- |
| Chiusi | Shallow lake | 11<br>(3 F, 6 FI, 2 D) | 8<br>(8 FI) | 8<br>(8 F) | 27<br>(18 F, 4 FI, 5 D) |  | 54 |
| Colfiorito | Wetland |  |  |  | 12<br>(6 F, 6D) |  | 12 |
| Fucecchio | Wetland |  |  |  | 12<br>(4 F, 8 D) |  | 12 |
| Massaciuccoli | Shallow lake,<br>wetland |  |  | 5<br>(5 F) | 12<br>(12 F) |  | 17 |
| Porta | Wetland |  |  |  | 5<br>(4 F, 1 D) |  | 5 |
| Trasimeno | Shallow lake |  |  | 10<br>(10 D) | 5<br>(5 D) |  | 15 |
| Vico | Deep lake |  |  |  | 16<br>(9 F, 7 D) | 6<br>(4 F, 2 D) | 22 |
|  | TOT | 11 | 8 | 23 | 89 | 6 | 137 |

### 2.7. Data Analyses

The temporal dynamics of spectro-functional traits across 2015-2020 years range were assessed by fitting an ordinary least squares linear regression line to spectral proxies time series and deriving for each spectral index the base value, as the intercept of regression line for year 2015 - WAVI_base_(2015-20), GLIbase(2015-20) NDSI_LMA_base_(2015-20) - and the overall change throughout the 5 years, as the difference between the intercept of regression line for year 2020 and the base value - ΔWAVI(2015-20), ΔGLI(2015-20) ΔNDSI_LMA_(2015-20).

The impact of meteo-climatic factors on *P. australis* in Central Italy was investigated by averaging the spectro-functional traits of all reed stands within each study area for a single year, separately for aquatic (flooded; N=25) and terrestrial (floating and dry; N=27) stands, and computing the reaction norms of WAVI, GLI and NDSI_LMA_ to Spring-Summer synoptic features (covering the 6 months spanning from March to August) derived from ERA5-Land model reanalysis data: average temperature (mean Temp. (Mar-Aug)), cumulated solar radiation (cum. Rad. (Mar-Aug)), and cumulated total evaporation (cum. Evap. (Mar-Aug)). The correlation between spectro-functional traits derived from Sentinel-2 data and meteo-climatic features was computed as Spearman’s *ρ* (with 95% confidence intervals and p-value).

Due to non-normality of samples – checked with Shapiro-Wilk test (p>0.05) – the variability of spectro-functional reed traits (and derived features) across ecological statuses (N=3: F, Fl, D), sites (N=7: Ch, Co, Fu, Po, Tr, Vi, Ma) or haplotypes (N=5: BS, K, L, M, VI) was tested with non-parametric Kruskal-Wallis One Way ANOVA, and *post-hoc* pairwise comparisons were successively performed via Dunn’s test, with p-value adjustment accounting for multiple comparisons computed with Benjamini-Hochberg method (FDR). The same statistical approach was used for assessing the differences among spectral-functional traits reaction to different meteo-climatic factors (temperature, radiation, evaporation quartiles).

The functional richness of each haplotype was calculated as the percentage of total trait space covered by each haplotype for each spectro-functional trait (Mason et al., 2005), separately for aquatic and terrestrial reed stands.

The partitioning of intraspecific variability in *P. australis* stands into components due to genotype (haplotype) or phenotypic plasticity was computed by fitting a linear mixed model (LMM) for each individual spectral proxy, including selected environmental factors as fixed effects (t_MarAug, rad_MarAug, ecol_stat, site) and haplotype as random effect.

All the analyses and graphing were performed using packages implemented in R v.4.0.3: i.e. *ggplot2* 3.1.0, *ggstatsplot* 0.7.2, *nlme* 3.1-152 (R Core Team, 2020).

## 3. Results

### 3.1. Haplotypes found

The combined alignment of trnT-trnL+rbcL-psaI regions used for the haplotype identification was 1627 bp length (Supplementary material S2). Four out of five haplotypes previously detected for the study areas – namely M, CHTR, CO and K (Coppi et al., 2018) - were confirmed in samples collected during 2020, and finally homologated following Saltonstall’s nomenclature (Supplementary materials S1). Haplotype VI was not resampled in 2020 because of logistic constraints. Table 2 provides an overview of haplotypes distribution among sampled stands and sites. The haplotype M was the most distributed and described for all sites. The CHTR haplotype was renamed as L, confirmed for lakes Chiusi and Trasimeno and described for the first time for Massaciuccoli wetland. The haplotype CO was defined as an intra-variation of the M haplotype (Supplementary materials S1) and was confirmed for Colfiorito wetland and Chiusi. We continued the analysis by including CO accessions within the M haplotype. The haplotype K was retrieved only for Chiusi, corroborating this haplotype as one of the less widespread and represented in the dataset. Lake Chiusi also hosts the new haplotype BS, which makes it the site featuring the higher level of haplotypic diversity (four haplotypes: L, K, M BS, and the so-called M variant CO) among those studied. The new haplotype BS differed from the haplotype M by the 16 bp insertion “AAAGTATTCTATAAAA” from the position 1494 of the intergenic spacer region rbcL-psaI (Supplementary materials S1).

### 3.2. Impact of meteo-climatic factors on spectro-functional traits of reed stands

Compared to the average values of the 1981-2010 period, in the last five years (2015-2020) the spring-summer meteo-climatic parameters (March-August) showed general increases of: average temperature (by 1.0-1.2 °C), cumulated incident solar radiation (by 66-105 MJ m^-2^), and cumulated total evaporation (by 1.8-6.6 mm of water equivalent).

While no consistent correlation was found between Spring-Summer meteo-climatic features and *P. australis* spectro-functional traits for terrestrial stands (growing on non-flooded terrain with different degrees of moisture), a negative response to average temperature was found for aquatic reed stands in terms of WAVI (*ρ*=-0.53, p=0.006) and GLI (*ρ* =-0.55, p=0.005), as well as positive response, yet slightly weaker, of NDSI_LMA_ to cumulated solar radiation (*ρ* =0.39, p=0.055), as shown in Figure 2.

**Figure 2.**
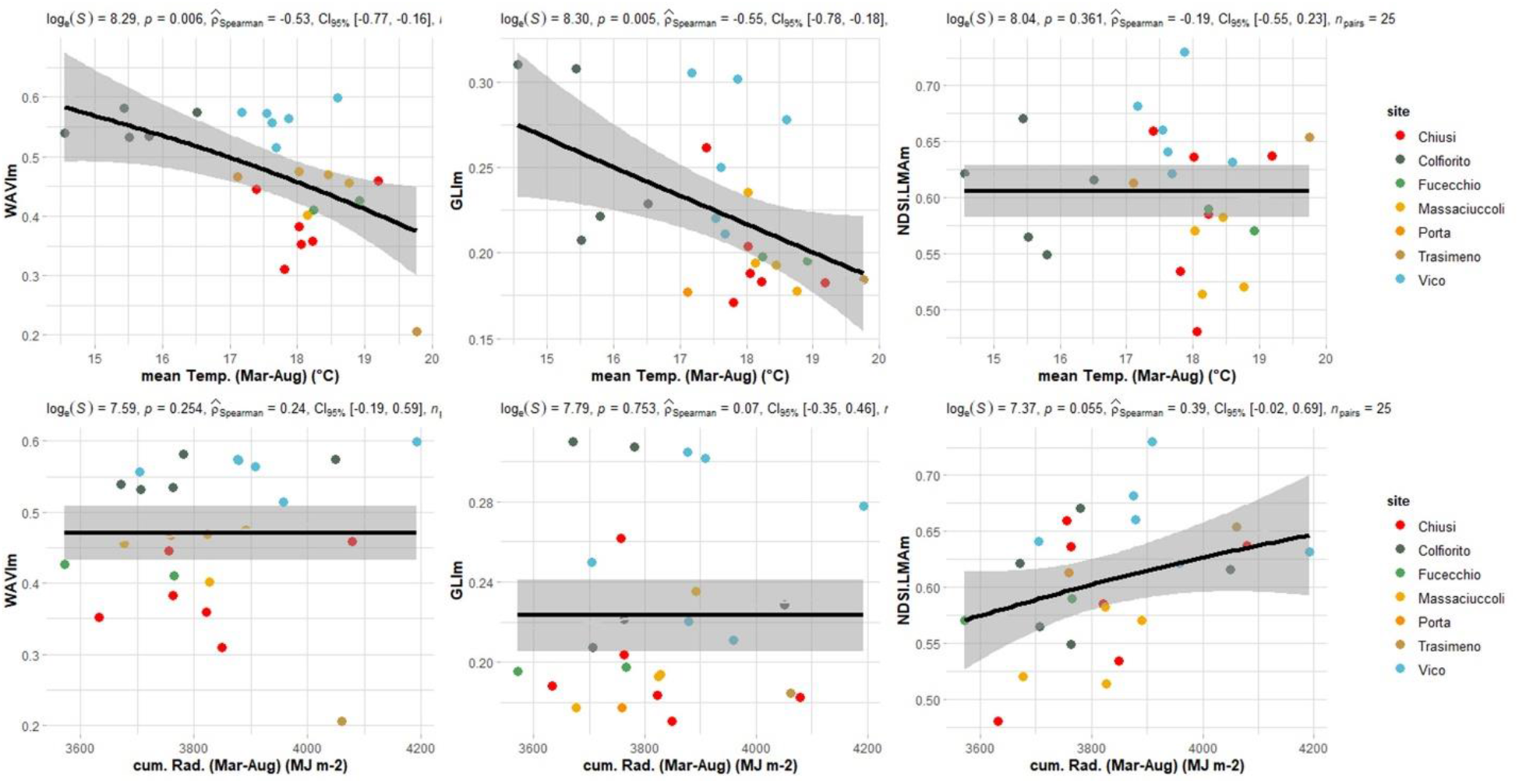
Reaction norms of spectro–functional traits (WAVI, GLI, NDSI_LMA_) derived from Sentinel–2 data to meteo– climatic features in Spring–Summer period (average temperature, cumulated solar radiation) for aquatic stands of *P. australis* in Central Italy, with superimposed generalized additive model (GAM) fitting (95% confidence intervals).

Specifically, spectral surrogates of canopy density (WAVI) and green leaf area (GLI) for aquatic reed stands tend to decrease with increasing temperature: in hotter seasons - i.e. when spring-summer temperature falls in the 1^st^ quartile of the 2015-2020 values (18.3-19.8 °C) - WAVI and GLI are lower on average by 19.0% (p=0.099) and 16.6% (p=0.078) respectively, than in the colder seasons - i.e. when spring-summer temperature falling in the 4^th^ quartile of the 2015-2020 values (14.6-17.1°C).

The spectral proxy of leaf dry biomass at canopy scale (NDSI_LMA_), responds positively to light availability: in the years that fall in the 1^st^ quartile of the 2015-2020 spring-summer cumulated values (3900-4192 MJ m^-2^), NDSI_LMA_ is on average 7.9% higher (p=0.028) than when solar radiation falls in the 4^th^ quartile of 2015-2020 spring-summer cumulated values (3572-3732 MJ m^-2^).

### 3.3. Intraspecific variability partitioning (haplotype vs. phenotype plasticity)

The analysis of variance partitioning in spectro-functional traits of *P. australis* stands, computed according to LMM using environmental factors as fixed effects (meteo-climate data, ecological status and site) and haplotype as random effect - summarized in Table 3 - highlighted that in general, variability due to haplotype strongly depends on the individual trait considered, ranging between 0% and 22% of total intraspecific variability across all reed stands. In aquatic stands, variability explained by haplotypes is higher for GLI and NDSI_LMA_, amounting to 22% and 34%, respectively. In terrestrial reed plots, comprising both dry and floating stands, instead, the variability explained by haplotypes is very low (< 2%), with virtually no effect of haplotype over the three traits considered.

**Table 3.**
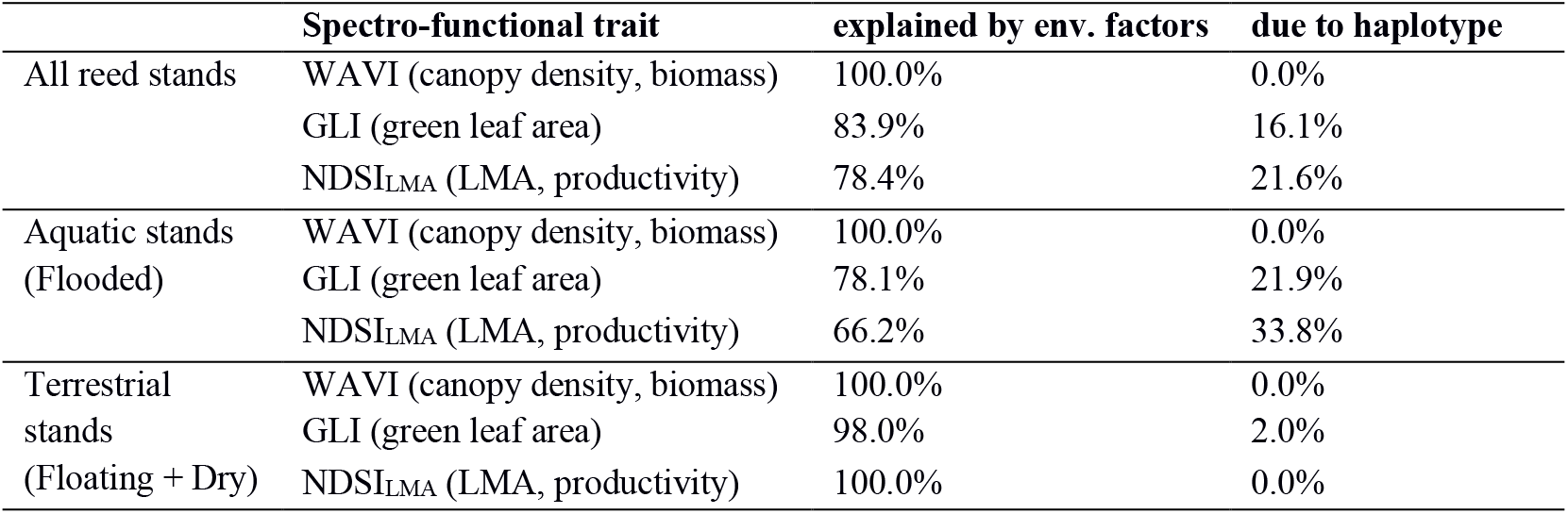
Partitioning of spectro-functional traits variance of reed stands.

|  | **Spectro-functional trait** | **explained by env. factors** | **due to haplotype** |
| --- | --- | --- | --- |
| All reed stands | WAVI (canopy density, biomass) | 100.0% | 0.0% |
|  | GLI (green leaf area) | 83.9% | 16.1% |
| | $\text{NDSI}_{\text{LMA}}$ (LMA, productivity) | 78.4% | 21.6% |
| Aquatic stands<br>(Flooded) | WAVI (canopy density, biomass) | 100.0% | 0.0% |
|  | GLI (green leaf area) | 78.1% | 21.9% |
| | $\text{NDSI}_{\text{LMA}}$ (LMA, productivity) | 66.2% | 33.8% |
| Terrestrial<br>stands<br>(Floating + Dry) | WAVI (canopy density, biomass) | 100.0% | 0.0% |
|  | GLI (green leaf area) | 98.0% | 2.0% |
| | $\text{NDSI}_{\text{LMA}}$ (LMA, productivity) | 100.0% | 0.0% |

### 3.4. Differences in spectro-functional traits of reed haplotypes

Figure 3 shows the distributions (as violin and box plots) and groups characterized by significant pairwise differences (p<0.05) of spectral functional proxies among *P. australis* haplotypes in our study areas, highlighting different patterns for aquatic and terrestrial stands (post-hoc test results in Supplementary materials (S3)).

**Figure 3.**
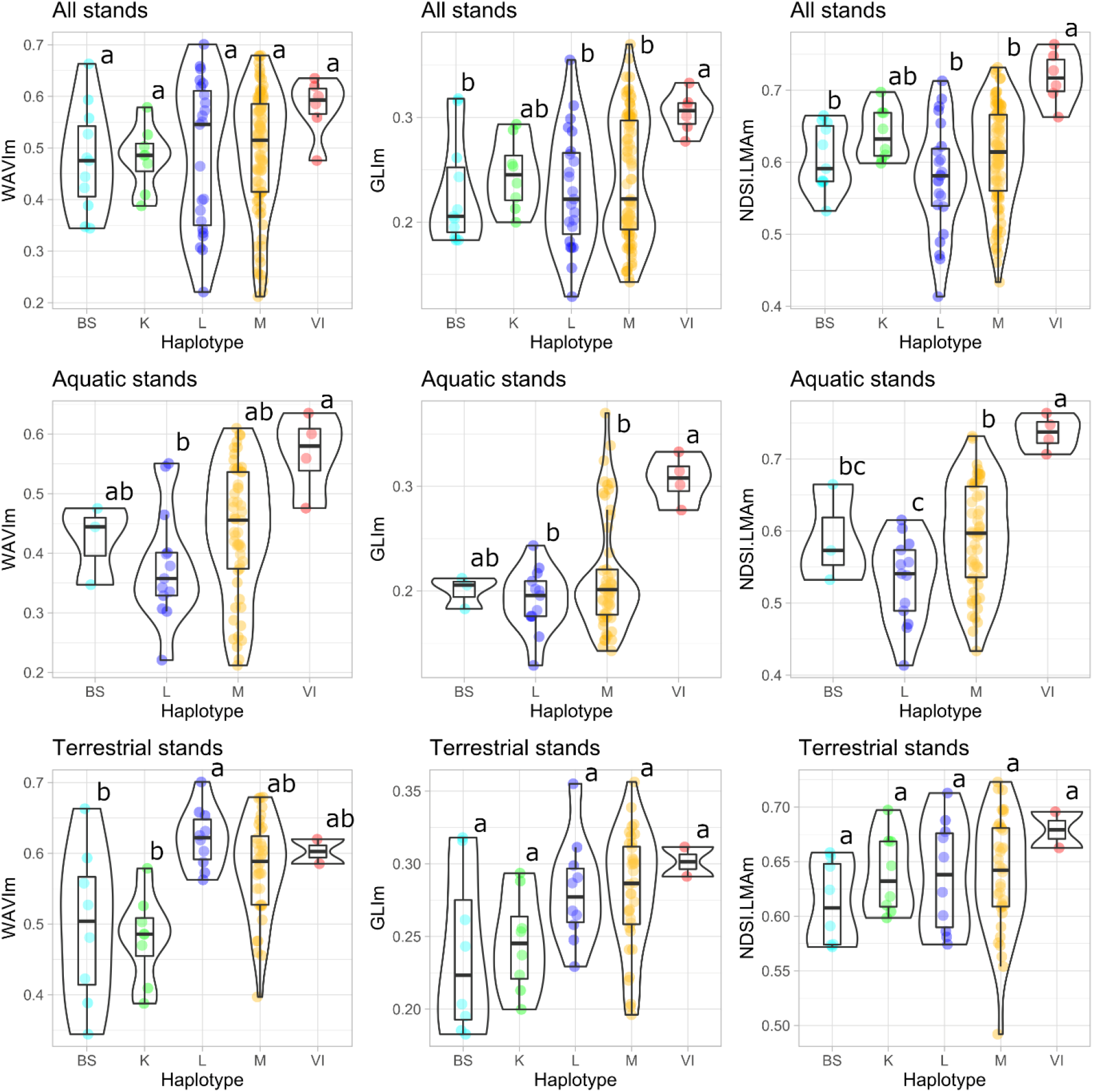
Violin plots (with encompassed box plots) showing range and distribution of spectro–functional traits (WAVI, GLI, NDSI_LMA_) across *P. australis* haplotypes in Central Italy: all stands pooled together (upper row); aquatic stands only (middle row); terrestrial stands only (lower row). Letters above the plots highlight groups characterized by significant differences (p<0.05) in pairwise haplotype comparisons, performed via Dunn’s post–hoc test (Benjamini–Hochberg adjustment).

Among haplotypes, L and M are the most variable in terms of spectro-functional traits, showing a pronounced functional plasticity. When all reed stands are considered, comprising stands growing in flooded to dry substrates, VI (median GLI=0.31) show higher GLI (p<0.03) compared to M, L and BS (median GLI=0.21- 0.22), and VI shows NDSI_LMA_ (median NDSI_LMA_=0.72) higher (p<0.005) than L, M and BS (median NDSI_LMA_=0.58-0.61).

When separated by ecological status, differences among haplotypes are generally less evident in terrestrial than in aquatic reed stands. Aquatic L stands score lower than homologous VI stands, in all spectral proxies (median relative difference>37%, p<0.02). Aquatic VI stands are characterized by higher scores of GLI and NDSI_LMA_ compared to stands of the most common M haplotype (median relative difference>23%, p<0.01), but not for WAVI.

Comparing aquatic and terrestrial stands of the most common haplotypes (L and M) shows that both score lower values of each spectro-functional canopy trait when growing in flooded conditions, more marked for WAVI and GLI (median relative difference>22%), than for NDSI_LMA_ (median relative difference<16%). While the GLI difference between aquatic and terrestrial stands tends to be similar for L and M (aquatic stands score 29% lower median GLI), the decrement in WAVI for aquatic stands of L is more marked than that of M (−42% vs −23% compared to terrestrial stands of the same haplotype, respectively).

Supplementary figure 1, as an example of the potential connected to the use of remote sensing data for deriving spatialized information about spectro-functional traits, portraits the patterns of relative variability in *P. australis* communities at within-site scale in Lake Chiusi (10 m spatial resolution), thus extending the information about local reed diversity beyond actually sampled points (marked with their respective haplotype on the map).

The breadth of the functional niche covered by different haplotypes is displayed by functional richness scores for the three spectro-functional traits considered, in Table 4.

**Table 4.** Functional richness (as percentage of total range) covered by each haplotype for each spectro-functional trait considered, separately for aquatic (flooded) and terrestrial (floating and dry) reed stands

| Spectro-functional trait | Ecological status | Haplotype |  |  |  |  |
| --- | --- | --- | --- | --- | --- | --- |
|  |  | BS | K | L | M | VI |
| WAVI<br>(canopy density, biomass) | Aquatic (Flooded) | 30.3% |  | 78.0% | 94.0% | 37.7% |
|  | Terrestrial | 89.4% | 53.5% | 38.8% | 79.2% | 9.7% |
|  | (Floating) | (63.8%) | (57.1%) |  | (60.7%) |  |
|  | (Dry) | (22.9%) |  | (45.5%) | (92.9%) | (11.4%) |
| GLI<br>(green leaf area) | Aquatic (Flooded) | 12.1% |  | 47.5% | 94.3% | 23.1% |
|  | Terrestrial | 77.9% | 54.0% | 72.5% | 92.3% | 11.7% |
|  | (Floating) | (56.6%) | (67.4%) |  | (58.1%) |  |
|  | (Dry) | (1.3%) |  | (78.5%) | (100.0%) | (12.7%) |
| NDSI<sub>LMA</sub><br>(LMA, productivity) | Aquatic (Flooded) | 37.8% |  | 57.6% | 85.2% | 16.3% |
|  | Terrestrial | 37.4% | 42.8% | 60.0% | 100.0% | 14.3% |
|  | (Floating) | (48.8%) | (65.4%) |  | (40.5%) |  |
|  | (Dry) | (1.5%) |  | (61.7%) | (100.0%) | (14.7%) |

The largest niche is occupied by the most common haplotype M, with functional richness >79%, followed by L, which cover >38% of total range across all traits, with BS covering a relatively wide niche (>37%) in terrestrial stands (including floating ones). Narrower niches are occupied by less spread haplotypes – i.e. VI found only in Lake Vico, K found only in Lake Chiusi – which cover <43% of trait range across haplotypes. Different patterns are specific each spectro-functional trait, particularly when reed stands are separated by their ecological status (aquatic vs. terrestrial stands).

### 3.5. Dynamics of spectro-functional traits of reed stands over the 2015-2020 period

Inter-seasonal dynamics of spectro-functional traits, summarized in Table 5, showed that floating reed stands (Fl) experienced a significant (p<0.05) decrement in WAVI (−0.031±0.003 y^-1^ for floating stands *vs*. −0.010±0.004 y^-1^ for flooded stands and −0.005±0.004 y^-1^ for dry stands) and NDSI_LMA_ (−0.019±0.003 y^-1^ for floating stands vs. −0.007±0.003 y^-1^ for dry stands). The trend is evident even when starting conditions in terms of trait scores – i.e. base value - are similar: WAVIbase(2015-20) scores 0.583±0.019 for floating stands, and 0.609±0.011 for dry stands; while NDSI_LMA_base_(2015-20) scores 0.688±0.009 for floating stands, and 0.649±0.010 for dry stands. This seems to be framed into the general environmental change occurring in Lake Chiusi – the only site where floating reed stands were found - during the last years (2015-2020). At the whole site scale, Chiusi clearly showed the most intense signs of reed canopy density loss, as surrogated by inter-annual WAVI change (Figure 4). Complementary to this, multitemporal maps of GLI and MDSI_LMA_ derived from Sentinel-2 clearly show the relatively fast dynamics of riparian plant communities in Lake Chiusi from 2015 to 2020 (Supplementary figures 2 and 3).

**Figure 4.**
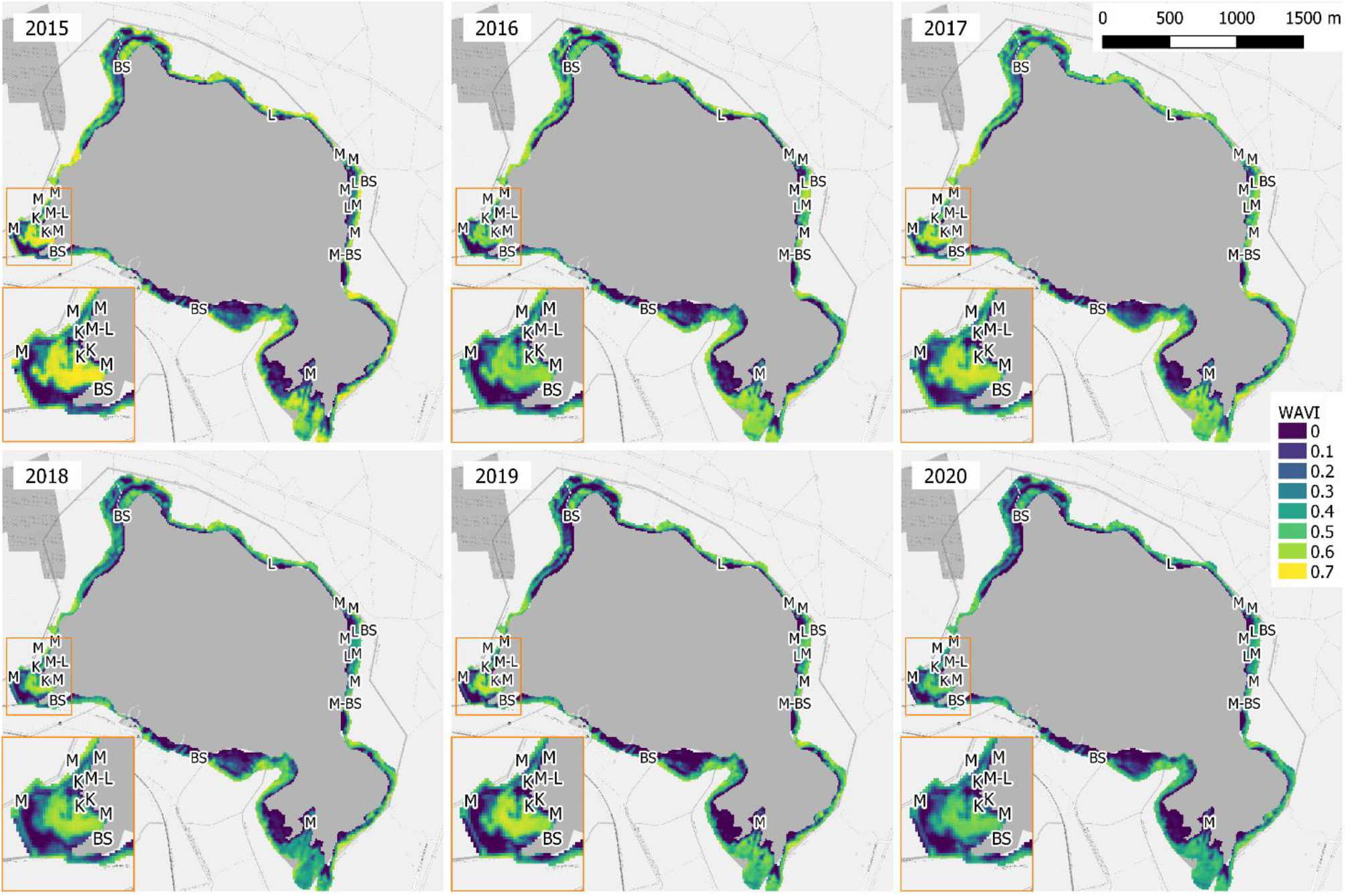
Multitemporal maps showing the temporal dynamics of WAVI (as a spectral proxy of canopy density) over *P. australis* communities of Lake Chiusi derived from Sentinel–2 scenes acquired at peak of growth (mid–July) from 2015 to 2020, with superimposed positions of genotyped reed stands (marked with haplotype code).

**Table 5.** Summary of significant pairwise differences (p<0.05) of *P. australis* spectro-functional traits dynamics indicators, i.e. base values and change throughout 5 years (2015-2020), categorized on the basis of stand ecological status, site and haplotype.

| Spectro-functional trait | Dynamics indicator | Ecological status (F, FI, D) | Site (Ch, Co, Fu, Tr, Vi, Ma)* | Haplotype (BS, K, L, M, VI) |
| --- | --- | --- | --- | --- |
| WAVI<br>(canopy density, biomass) | ΔWAVI(2015-20) | FI < (F, D) | Fu > (Ch, Ma, Tr)<br>Vi > Ch | BS < (M, VI)<br>K < (M, VI) |
|  | WAVI<sub>base</sub>(2015-20) | D > F | Tr > (Ch, Ma) | No differences |
| GLI<br>(green leaf area) | ΔGLI(2015-20) | No differences | Fu > (Ch, Co, Tr, Vi) | No differences |
|  | GLI<sub>base</sub>(2015-20) | D > F | Ma < (Co, Tr, Vi) | No differences |
| NDSI<sub>LMA</sub><br>(LMA, productivity) | ΔNDSI<sub>LMA</sub>(2015-20) | FI < D | Ch < (Fu, Vi) | No differences |
|  | NDSI<sub>LMA_base</sub>(2015-20) | FI > (D, F) | Ma < (Ch, Vi) | K > (L, M)<br>VI > (L, M) |

\* Porta (Po) excluded from this analysis because satellite data cover only 1 year (2019)

ΔWAVI(2015-20) amounts to −0.024±0.003 y^-1^ in Chiusi (significantly different, p<0.001, from Fucecchio and Vico) compared to −0.010±0.005 y^-1^ in Colfiorito (p=0.06), and −0.008±0.005 y^-1^ in Trasimeno (p=0.09); Massaciuccoli is the only site where of reed density loss through time is not much different from Chiusi (−0.015±0.012 y^-1^, p=0.36). Reed stands conditions did not experience a statistically significant decrement of reed spectro-functional traits in 2015-2020, only in Fucecchio and Vico.

Less spread haplotypes (K, VI) are mainly separated from more common haplotypes (L, M) by baseline NDSI_LMA_ scores, which is higher in K and VI stands (p<0.05).

## 4. Discussion

Competition among plants can occur along different spatial and temporal axes: along the growth cycle, it unfolds both in horizontal and vertical directions, affecting functional traits and the adaptive trade-offs of interacting individuals (Bittebiere et al. 2017 and references therein). Following this framework, interaction among haplotypes in *P. australis* communities would involve variations of traits linked with the competition for light capture and use for optimising the photosynthetic efficiency (vertical growth, canopy density) and/or traits involved in gain of supremacy in space occupation (horizontal spreading of rhizomes). From a temporal dynamics point of view, competitive exclusion in common reed stands may be delayed by intraclonal aggregation in the short term (Silvertown et al., 1992), but clonal fragmentation may occur over medium to long time scales. The life span of *P. australis* rhizomes seem to decay in a period of around 3–7 years (Silvertown et al., 1992), leaving space to different genetic integrated units which might compete vigorously against each other because of their converged functional traits constitution. Five out of the seven sites we investigated in the present study, Fucecchio and Porta being the exceptions, showed the simultaneous occurrence of more than one haplotype, with signs of dynamic interaction among genotypes that produces a fine-scale mosaic of functional variability of local *P. australis* stands. Moreover, in the same areas several reed stands have experienced marked change in their extension - mostly shrinking - in the last 25 years (Lastrucci et al., 2017; Gigante et al., 2017). Thus, we hypothesised that narrowly adapted clones have selected optimal microhabitat during the last years thanks to their functional trait plasticity. Indeed, previous research on the same species (Hu et al., 2015) has evidenced significant amount of variation in leaf economic traits across and intra-sites. In a time-oriented point of view, during the last growing seasons (2015-2020), some key meteo-climatic factors showed slight by significant increment compared to the average of 1981-2010 period across the study areas, in terms of mean temperature (by 1.0-1.2 °C), cumulated incident solar radiation (by 66-105 MJ m^-2^) and cumulated total evaporation (by 1.8-6.6 mm of water equivalent). These variations were mirrored by changes in spectro-functional traits of *P. australis*, which showed a significant reduction in GLI in the hotter years. GLI can be considered as a spectral proxy for Leaf Area Index (LAI), which represents a key-variable parametrising productivity and growth at canopy scale, also linked to evapotranspiration capacity (Burba & Verma, 2001). Green LAI tend to respond to variation in temperature very differently from species to species (Iio et al., 2014), but negative association for *P. australis* has been observed by Anda et al. (2017) that have measured lower LAI scores for stands growing in dryland conditions, prone to periodic water stress. Contradictory results in morphological signatures were nevertheless provided for common reed stands growing in different water depth condition (Engloner et al., 2009). Indeed, our dataset showed values of GLI significantly higher for dry than for flooded stands, indicating a different response of two ecological statuses to temperature variation.

Similarly to what was previously found across many other plant species (Poorter et al., 2010), the studied reed stands also respond to light availability showing an higher spectral proxy for LMA (NDSI_LMA_) with increasing cumulated radiation, indicating a general rise in thickening of leaf blade or a denser tissue thereof, or both (Wright et al., 2004). Nevertheless, NDSI_LMA_ dynamics among common reeds stands exhibited different patters depending on reed ecological status, site location and haplotype. In particular, floating island stands (Fl) evidenced a significant reduction of NDSI_LMA_ and WAVI values, suggesting a decrease in biomass production and culms density of the stands during the last years (2015-2020). The finding suggested a general time-dependent deterioration in the state of the reeds in this site in recent years, compared to the work of Gigante et al. (2014), which described the floating reeds on Lake Chiusi as healthy and not susceptible to decay from data collected in summer 2011. Based on our framework, the time-dependant-reduction of the floating island reed stands could be indirectly related to haplotype composition: indeed, Chiusi is the only site that host floating stands, located in the northern and western part of the lake. Three out of five haplotypes were identified for this ecological category (M, BS and K), and two of these haplotypes (BS, K) showed a very reduced distribution. It is conceivable that under the pressure of environmental change in Lake Chiusi, those haplotypes have experienced intense competition and have narrowly adapted to the microhabitat constituted by floating condition. On the other hand, they may have become more sensitive to changes in local impact factors, such as wave movement. One possible interpretation of this finding is that the extensive reduction of common reed area in Lake Chiusi during the last 3 decades (Gigante et al., 2014; Lastrucci et al., 2019) could have reduced the protection of the open water from the wind, increasing the wave movement events and consequently their incidence on remnant reed stands (pers. obs.).

The dualism we observed between more common haplotypes and less spread ones across our study areas suggests the coexistence of two main evolutionary strategies deployed by this species: maximize phenotypic plasticity, which may favour colonizing new habitats (high functional richness, even with generally lower canopy traits implying suboptimal productivity, such as for stands with haplotypes L and M), or maximizing productivity through specialization to peculiar site conditions (low functional richness, but higher scores of spectral proxies for traits connected to canopy density/biomass, and eventually productivity, such as for stands with haplotype VI).

Overall, our results showed that haplotypes explained a tangible portion of total intraspecific variability in spectral proxies of canopy traits across sampled reed communities, of variable magnitude (0-34%), depending on ecological status and/or spectro-functional trait considered. The higher portion of variability due to haplotype in aquatic stands could be linked to more intense selective pressure exerted on reeds by rhizome flooding, that cannot be tackled only by phenotypic plasticity (Engloner et al., 2010; Gigante et al., 2011, 2014; Lastrucci et al., 2016a, 2017)

The most common haplotypes (L and M) showed significantly lower values of each proxy under flooded than in dry conditions, of major entity for WAVI and GLI, and minor one for NDSI_LMA_. Moreover, while the difference between flooded and dry stands for GLI was similar for L and M, flooded L stands scored a markedly larger decrement in WAVI compared to terrestrial ones. Such a difference suggests that canopy biomass of aquatic L stands tends to be lower than that of other haplotypes, and this could be putatively connected to the higher impact of dieback on flooded L stands. This outcome is in line with previous works that showed macro-morphological evidence, such as clumping habitus or lower culms diameter, commonly described for flooded reeds as generally associated with the dieback syndrome of *P. australis* (Lastrucci et al., 2016c; Coppi et al., 2018). Related findings may help shedding some light on drivers of common reed dieback in European aquatic stands, a largely investigated yet to date still poorly understood phenomenon, in terms of cause-effect processes (Ostendorp, 1989; Van der Putten, 1997; Gigante et al., 2011; Gigante et al, 2013; Gigante et al., 2014; Coppi et al., 2018).

## 5. Conclusion

Our results dealing with reaction norms of spectro-functional traits to macro-scale meteo-climatic factors bear implications with respect to ongoing scenarios of climate change and in particular summer warming, as trends of increasing temperature appear to be related to decreased canopy density (WAVI) and green leaf area (GLI) in flooded reed stands. This outcome appears particularly interesting and indicates the advisability of further investigations in this direction to understand better the processes underpinning common reed decline in Europe, and in particular the Mediterranean region.

Taking advantage of medium-high resolution satellite data collected by Sentinel-2 constellation, freely available since summer 2015 and now granting an observation frequency of up to 5 days over the majority of globe, we have investigated the canopy reflectance diversity in the dominant macrophyte *P. australis*, its relation to the haplotype diversity, the meteo-climatic parameters, and the different ecological condition of reed stands. Multispectral satellite data covering different years and systems have allowed exploring intraspecific variations in spectro-functional traits of *P. australis* within and among sites at spatial scales virtually impossible to assess from in situ survey only. Through this, we enlarged the knowledge on the distribution in Central Italy of previously known *Phragmites australis* haplotypes, as well as to identify a new haplotype (BS). Furthermore, we provided new evidence on haplotype-haplotype and haplotype-by-environment interactions for this species. In particular, the interaction among haplotypes and its relation to the spectro-functional variability of *P. australis* stands.

## Acknowledgements

This work has been supported by the project “macroDIVERSITY”, funded by the Ministry of Education, University and Research, PRIN 2017 [grant 2017CTH94H]. The authors would like to thank M. Meloni for their help during the sampling.

## Statements and Declarations

The authors exclude any potential sources of interest directly or indirectly related to the work submitted for publication.

## Data Availability Statement

The original sequences produced for the study have been deposited into the database GenBank: www.ncbi.nlm.nih.gov/genbank

The spectro-functional traits data are available from the authors upon reasonable request.

